# Predictions of Skin Permeability Using Molecular Dynamics Simulation from Two-Dimensional Sampling of Spatial and Alchemical Perturbation Reaction Coordinates

**DOI:** 10.1101/2022.02.10.479880

**Authors:** Magnus Lundborg, Christian Wennberg, Jack Lidmar, Berk Hess, Erik Lindahl, Lars Norlén

## Abstract

A molecular level understanding of skin permeation may rationalize and streamline product development, and improve quality and control, of transdermal and topical drug delivery systems. It may also facilitate toxicity and safety assessment of cosmetics, skin care products. Here, we present new molecular dynamics simulation approaches that make it possible to efficiently sample the free energy and local diffusion coefficient across the skin’s barrier structure and predict skin permeability and the effects of chemical penetration enhancers. In particular, we introduce a new approach to use two-dimensional reaction coordinates with so-called Accelerated Weight Histograms, where we combine sampling along spatial coordinates with an alchemical perturbation virtual coordinate. We present predicted properties for twenty permeants, and demonstrate how our approach improves correlation with *ex vivo/in vitro* skin permeation data. For the compounds included in this study, the ob-tained 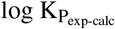 mean square difference was 0.8 cm^2^ h^−2^.

## Introduction

Transdermal and topical drug delivery allows for non-invasive, pain-free, continuous drug administration with reduced side effects and increased patient compliance compared to per oral or intravenous drug delivery.

Assessment of skin permeability is important for the design of percutaneous drug delivery systems. Today, the dominating means of investigating drug skin permeability and the effects thereon of different delivery vehicle excipients is by *ex vivo/in vitro* testing. There are, however, ethical problems involved in the study of percutaneous drug delivery using excised human or animal skin, or using animals *in vivo*.(1, 2) There are also issues of the translation of animal drug delivery data to humans.(1–3) When using human skin, there are complications of inter- and intraindividual variation as well as of inter-laboratory differences.(1, 4) Furthermore, *ex vivo/in vitro* testing yields no information about how compounds permeate skin.

*In silico* modeling is an interesting alternative to predict skin permeability and to gain insights into how compounds are absorbed in skin (5), but it requires both faithful models of the skin system and methods that can sample the process efficiently. A skin permeation prediction model relevant for human skin, encompassing the effects of penetration enhancers and other drug delivery vehicle excipients, could help improve drug delivery formulation design.

We have previously shown that an atomistic model of the skin’s barrier structure, i.e., the intercellular lipid matrix of the stratum corneum, validated against cryo-electron microscopy (cryo-EM) data from near-native skin (6) (Fig. 1), could be used in molecular dynamics (MD) simulations to predict the permeability of molecules through skin (7). These models made it possible to reproduce the effects of chemical penetration enhancers on the skin’s barrier structure at least qualitatively (7), which enabled a molecular-level understanding of their mechanisms of action, but the amount of sampling required and limited convergence made it difficult to accurately match experimental values. Improving the efficiency of sampling for this type of complex system has led to a number of improvements of simulation protocols, in particular with respect to the sampling of the free energy landscape and local diffusion coefficient of drugs passing through the skin’s barrier structure (8).

**Fig. 1.**
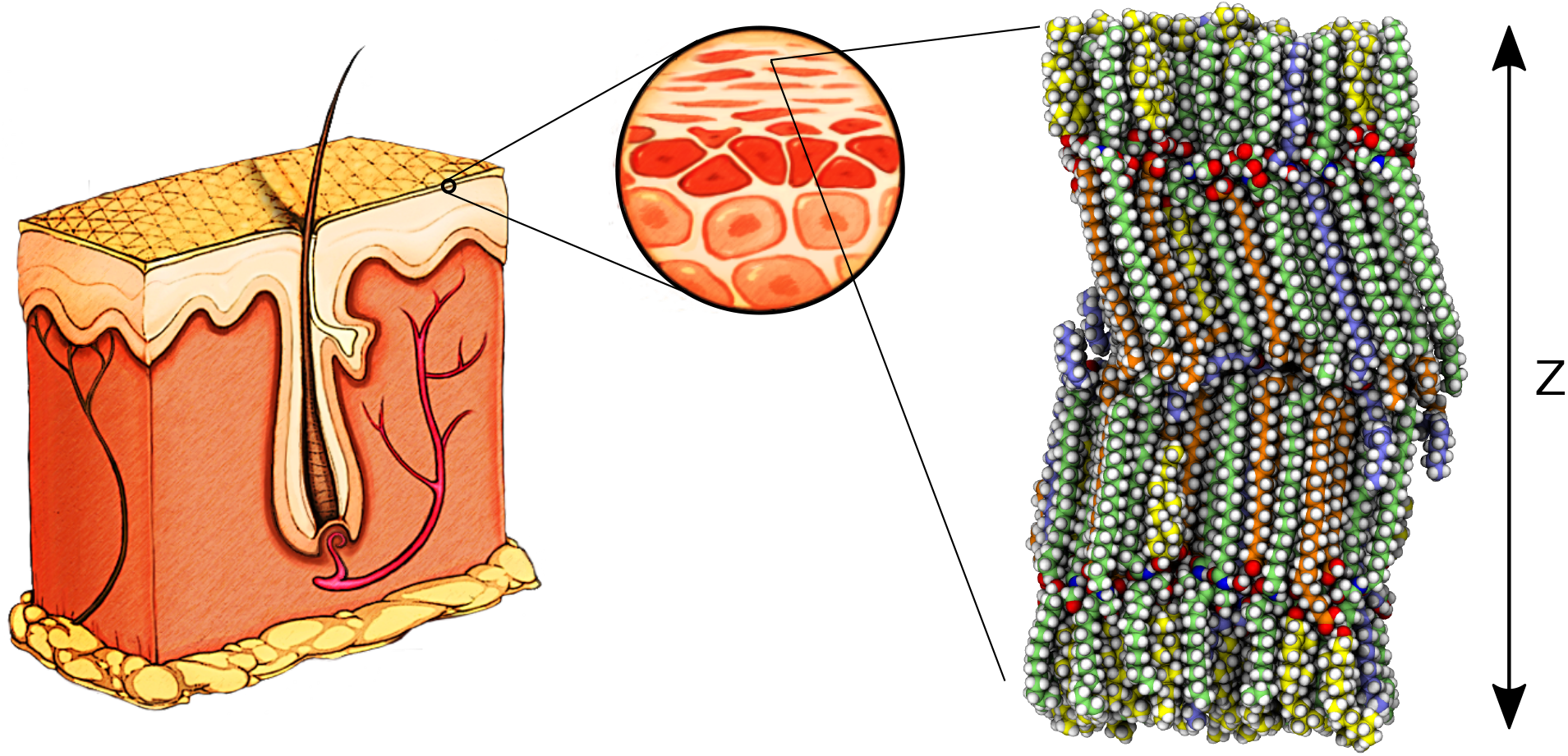
A schematic representation of the structure of epidermis and dermis. To the right is shown a snapshot of the atomistic model of the intercellular lipid structure of human stratum corneum that constitutes the skin’s main permeability barrier.(6) The arrow indicates the main permeability direction, also referred to as the *Z* dimension. In the MD simulations the system is in practice repeated infinitely in the three dimensions. The carbon atoms are coloured based on the molecule type, where ceramides are green, acyl ceramides (ceramide EOS) are light blue, free fatty acids are orange and cholesterols are yellow. Hydrogen, nitrogen and oxygen atoms are colored white, blue and red, respectively.

We here apply the Accelerated Weight Histogram (AWH) method, using reaction coordinates that combine a spatial dimension, pulling the molecule across the barrier structure, with a second alchemical free energy dimension. (8) This has made it possible to sample the free energy landscape and local diffusion coefficient of 20 chemical compounds with molar masses ranging from 18 gmol^−1^ to ∼300 gmol^−1^ and logP_o/w_ from −2.1 to 4.6. It is expected that such sampling, allowing decoupling of the permeant, will be efficient (9) in systems with long correlation times, such as the skin’s barrier structure. An improved agreement between calculated and measured permeability coefficients is presented. It is shown that, using the AWH method, *ex vivo/in vitro* measured skin permeability data can be accurately predicted, even though the *in silico* model selectively accounts for the skin’s barrier structure and not for the complete skin.

### Molecular dynamics skin permeation modeling

Quantitative Structure-Permeability Relationship (QSPR) methods (10–12) are commonly used to predict skin permeability coefficients. Molecular dynamics (MD) simulations constitute an alternative and/or complement to QSPR methods. MD simulations are significantly slower than QSPR methods, but not as limited to a specific applicability domain. Further, incorporating the effect of formulation excipients on the permeability of active pharmaceutical ingredients (APIs) requires retraining of QSPR models, preferably for each combination of excipients. There have, nevertheless, been QSAR (Quantitative Structure-Activity Relationship) (13) studies involving chemical permeation enhancers, but information about these excipients’ mechanisms of action has been difficult to obtain. With MD simulations, formulation excipients can be incorporated in the skin’s barrier structure in their most probable concentrations, locations and orientations, based on their local free energy potentials. This enables studying how they interact with the barrier and how they affect the permeability of one or more APIs.

One limitation of the MD simulation approach presented here is that only the structure of the main permeability barrier in skin (14, 15), and not the complete skin, is studied in detail. However, it would be possible to complement the calculated permeability coefficients with mathematical models accounting for corneocyte permeability as well as for diffusion through the viable epidermis and dermis.(16, 17)

In our previous studies we showed that it is possible to predict the permeability coefficient of different chemical compounds through an atomistic model of the skin’s barrier structure (6, 7), illustrated in Fig. 1. Therein we used MD simulations to pull the permeants through the barrier system using a stiff spring and the forward-reverse (FR) method to calculate the permeability coefficients.(18–21)

Based on our previous proof-of-concept study (7) we have studied the effects of the pulling speed on the calculated permeability coefficients in more detail. In the supplementary material (Fig. S1) we illustrate sampling problems using potentials of mean force (PMFs), i.e., the relative free energy difference across the system, of testosterone that we have experienced to be a permeant that needs careful sampling for obtaining accurate results. The general observation was that with slower pulling speeds the PMFs grew more detailed and the result free energy barriers became lower. We attribute this to the skin’s barrier lipid system being in a gel-like state and that the permeant needs to be pulled slowly to remain in a state near equilibrium, even if forward-reverse pulling is a nonequilibrium method (20). The experience that slower pulling speeds resulted in lower PMFs correlated in time with the publication by Wang and Klauda (22), in which they stated that all 80 (approximately) stacked lipid bilayers of the skin’s barrier structure must be taken into account (22–24), since the calculated permeability coefficient is not actually an average speed through the system, even if its unit would indicate that. The effect is that the permeability coefficient should be divided by 80, or 1.9 be subtracted from its log value. When taking 80 stacked lipid bilayers layers into account for calculating the permeability coefficients using the data from our previous study(7) the 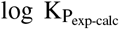 mean square difference was approximately ∼11 cm^2^ h^−2^ (cf. ∼3 cm^2^ h^−2^ in the original work(7)). This made us conclude that very slow pulling speeds might be required to obtain correct PMFs. This would, in turn, require very long simulation times for each pulling simulation (see Figs. S1 and S3) - potentially so long that entirely different sampling approaches are required to predict permeability with reasonable amounts of computing time. Since it is not the main topic of this paper we refer to the supplementary information for more discussions about FR pulling and umbrella simulations.

A permeant’s skin permeability coefficient is mainly determined by the height of the peaks, relative to the depth of the troughs, of its potential of mean force (PMF), i.e., the local free energy across the skin’s barrier structure, assuming that the passage through the barrier structure is the rate limiting step for diffusion. On the other hand, the partitioning of a permeant from a formulation into the skin’s barrier structure is primarily depending on the depth of the troughs of the permeant’s PMF. It is therefore important that the MD simulations can reproduce the whole PMF as accurately as possible, given the available computational resources.

### Skin permeability calculations with the Accelerated Weight Histogram method

The Accelerated Weight Histogram (AWH) method (25, 26) is an extended ensemble technique, in which an adaptive bias is used to flatten the free energy landscape. The applied bias enables sampling high free energy (low probability) configurations to the same extent as low free energy states. It is also possible to customize the target sample distribution to focus more on regions of the free energy landscape that are of higher interest.

In the work presented here we have used the AWH method to improve the sampling of the permeability of 20 permeants through the skin’s barrier system. Since GROMACS 2021 it is possible to use a two-dimensional AWH sampling with an alchemical free energy dimension (8) combined with a spatial dimension, i.e., pulling the permeant across the skin’s barrier system. The alchemical free energy calculations imply that the interactions of the permeant with its surroundings can be gradually decoupled (switched on or off), thereby estimating the free energy of insertion. This means that there is no need for separate calibration of the PMFs in relation to the delivery vehicle, as all points along the PMF will be calibrated compared to a vacuum state, in the same way as the solvation free energies are calculated. This also improves the sampling in regions with very slow diffusion (long correlation times), since turning off the interactions with the surroundings will let the permeant leave that region in order to sample other parts of the landscape, and later return in a different configuration.(9) Fig. 2 shows a general overview of the method applied to calculating the PMF of testosterone through the skin’s barrier structure. With the AWH method it is also trivial to extend the simulations until they are sufficiently converged. This solves the primary sampling problem from previous studies, and it also enables us to calculate the local diffusion coefficient based on the AWH friction metric along the spatial dimension. The free energy of solvation in the solvent, or formulation, should still be calculated, which can be done in a separate one-dimensional alchemical AWH simulation.(8)

**Fig. 2.**
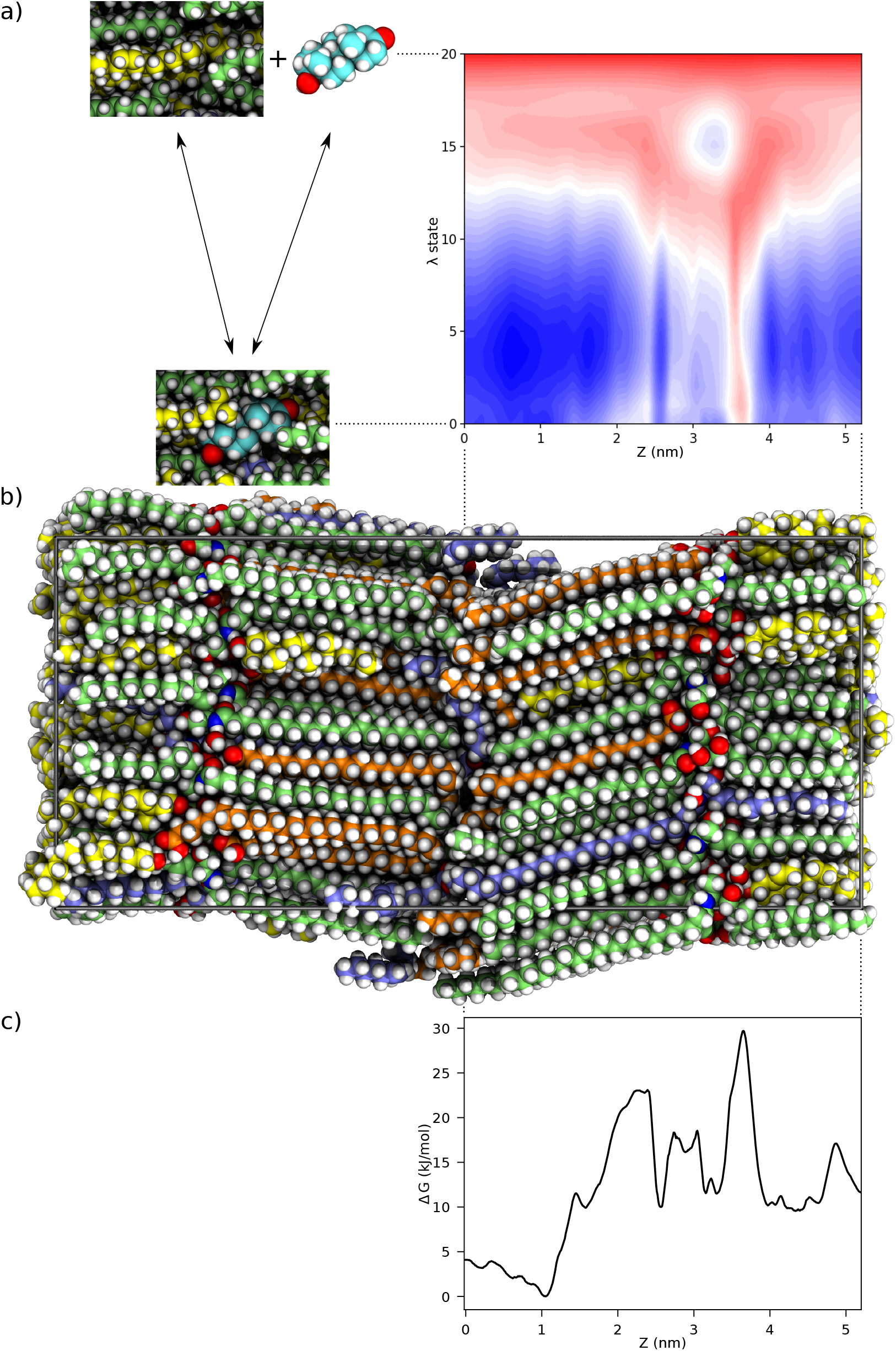
Calculating the PMF of testosterone through the skin’s barrier structure. a) on the right, shows the two-dimensional free energy landscape of testosterone where the alchemical free energy *λ* state is shown on the *Y* axis — 0 is fully interacting and 20 is fully decoupled (as illustrated by the miniatures to the left of the free energy landscape). On the *X* axis the *Z* coordinate (in nm) through the simulated system is shown. b) shows the skin’s barrier system, aligned to illustrate how the PMF is mirrored around 0 (the middle of the system). c) shows the PMF of testosterone through the skin’s barrier structure of the fully interacting *λ* state (0 on the *Y* axis in a).

### Calculating permeability coefficients as a two-step process

In simulation systems where the delivery vehicle, or solvent, is present along with the permeability barrier it is common to calibrate the PMF by setting it to zero in the delivery vehicle.(27) If the delivery vehicle is not simulated, like in the case of the stacked bilayer system studied here, the PMF can be calibrated based on a separate solvation free energy calculation.(7) However, once the permeant has partitioned from water into the barrier structure it is the relative free energy barriers that are the important factor, with troughs in the PMF acting as traps. Shifting the PMF so that its minimum is 0 is similar to what Wang and Klauda (22), Venable et al. (28) have done before.

In this project we have used a two-step process for the analysis of the permeability. The PMF, for calculating the permeation resistance, of one layer was calibrated relative to the hydration free energy. The permeation resistance through the remaining 79 layers was calculated with the PMF minimum set to 0.

Applying this two-step permeability process, as well as accounting for the 80 stacked bilayers (described above), changed the log 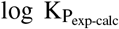 mean square deviation in our proof-of-concept study (7) from ∼3 cm^2^ h^−2^ (using the reference data in the original article(7)) to ∼17 cm^2^ h^−2^. It should be noted that the reference data used in that study was not corrected to account for the neutral species of ionizable compounds (codeine, naproxen and nicotine). If doing that, the log 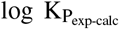 mean square deviation would have been even larger (∼19 cm^2^ h^−2^).

### Alchemical transformations in a gel phase system

It is widely accepted that it is more efficient to perform electrostatic and Lennnard-Jones transformations (coupling/decoupling) in sequence to ensure that the Coulomb interactions are switched off before Lennard-Jones interactions are and vice versa.(29) This avoids overlap of atoms when the attractive Coulomb forces are strong and the repulsive Lennard-Jones forces are turned off or are very weak. However, the skin’s barrier structure is in a gel-like state with very low mobility. We have found that the optimal orientation of a permeant molecule might be completely different in a state with its Lennard-Jones interactions turned on and Coulomb interactions turned off, compared to in its fully interacting state, and that the interconversion between the orientations can take very long when the Lennard-Jones interactions are turned on. For example, consider inserting (turning on interactions of) 1-decanol in the sphingoid chains of the skin’s barrier structure. If first turning on Lennard-Jones interactions the probability of inserting the molecule with the hydroxyl group oriented away from the head group region, into the interface between the lipid chains, will be approximately the same as inserting it in its most favorable position with the hydroxyl group oriented towards the headgroup region. When turning on the Coulomb interactions it will be unfavourable if the molecule is oriented in the opposite direction and it will take very long for it to turn around. If turning on the Lennard-Jones and the Coulomb interactions simultaneously the electrostatic interactions will favour inserting the molecule correctly from the start.

In this project we decided to use the same, simultaneous interaction transformation, settings for both the two-dimensional alchemical free energy/spatial AWH simulations as well as for the one-dimensional AWH alchemical hydration free energy calculations, even if the efficiency might not be optimal for the latter. It should be noted that this problem is at least partially avoided if the alchemical free energy simulations are started from a fully interacting and equilibrated state, or as mentioned above, in a system with fast diffusion in all dimensions.

## Methods

The atomistic model of the human skin’s barrier structure presented by Lundborg et al. (6), and originally called 33/33/33/75/5/0.3^1^, was used as the starting structure for the simulations. The starting model had been equilibrated for approximately 270 ns, 250 ns of which were without restraints (6).

All MD simulations in this study (not including the data presented in Figures S1 and S2) were performed using GROMACS 2021 and the second beta version of GRO-MACS 2022 (30–32), using the Accelerated Weight Histogram (AWH) method for alchemical free energy calculations (8, 25, 26). A source code modification enabled symmetrizing AWH sampling along a spatial (pulling) reaction coordinate dimension and also changed the number of AWH blocks for autocorrelation analysis from 64 to 128. These changes are available from the GROMACS gitlab repository (33).

Van der Waals interactions had a cutoff of 1.2 nm with a smooth force-switch from 1.0 nm to 1.2 nm. The simulations were run without a dispersion correction for energy and pressure. Coulomb interactions were calculated using PME (34) with a radius of 1.2 nm. Bonds to hydrogen atoms were constrained using the P-LINCS algorithm (35, 36). TIP3P (37) parameters were used for water molecules. For the lipid molecules the CHARMM36 lipid force field (38, 39) was used. Ceramide parameters were modified to more accurately reproduce the ceramide NP crystal structure.(40) as described in Ref. 6. In order to allow a 3 fs integration time step, hydrogen atoms were made three times heavier by repartitioning the corresponding mass from their bound heavy atoms. Temperature was set to 305.15 K by using a stochastic dynamics integrator (also referred to as a velocity Langevin dynamics integrator) with a time step of 3 fs and with a time constant *τ* of 2 ps (corresponding to a friction constant of 0.5 ps^−1^). The pressure was set to 1 atm and controlled using a stochastic cell rescaling barostat (41) with a time constant of 1.0 ps and a compressibility of 4.5 × 10^−5^ bar^−1^. In the skin’s barrier system the pressure coupling was semi-isotropic with no compressibility in the *Z* dimension. For all permeants, at least one simulation was performed with unmodified hydrogen masses and a 2 fs time step in order to verify that the longer (3 fs) time step did not affect the results.

Along the alchemical free energy dimension, 21 equidistantly distributed *λ* states were used for decoupling both van der Waals and Coulomb interactions simultaneously. There were 10 or 100 steps between each sampling of the alchemical free energy *λ*-value and 10 samples per update of the *f*_*λ*_. There was no observable difference between sampling every 10 or 100 steps, but the simulations were faster when sampling less frequently. The estimated initial error was set to 10 kJ mol^−1^. Soft-core transformations (42) with *α* = 0.5 and *σ* = 0.3 nm were applied to both the van der Waals and Coulomb interactions of the solute.

Topologies, i.e. inter- and intramolecular interaction parameters, for all permeants and formulation components except water, were generated using STaGE (43), which in turn uses Open Babel (44) and MATCH (45) to generate GROMACS topologies.

All images representing molecules were prepared using Tachyon (46) in VMD (47).

### Hydration Free Energy Calculations

The hydration free energy was calculated starting with a water box large enough to solvate the molecule. The molecule was inserted into the system at a random position with its interactions to the surroundings turned off, as if in vacuum. AWH was used to sample the alchemical free energy *λ* states (8) to calculate the solvation free energy. The initial AWH histogram size, which determines the initial update size of the free energy, was set indirectly by specifying an estimate of the diffusion constant along the alchemical free energy *λ*-axis in combination with a rough estimate of the initial error. An input diffusion constant of 1 × 10^−3^ ps^−1^ was used for the hydration free energy calculations, which means that it is estimated to take approximately 1 ns to cross the alchemical dimension for one walker. The initial error was set to 10 kJ mol^−1^. 16 communicating walkers were run in parallel, with the requirement that only simulations that covered the whole alchemical reaction coordinate counted towards the covering check in the initial AWH stage. The simulations were 60 ns long per walker for a total simulation time of 960 ns per solute. For a more detailed description of alchemical hydration free energy calculations using AWH see Lundborg et al. (8).

### Permeability Coefficient Calculations

The permeability coefficient, *K*_*P*_, and the permeation resistance, *R*, were calculated as follows(48):

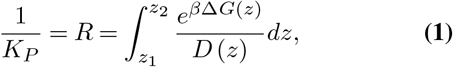

where Δ*G* is the difference in free energy, compared to the reference state, and *D* is the local diffusion coefficient, across the skin’s barrier structure (*dz*).

As mentioned above, we assumed a two-step process for calculating the permeability coefficient. The PMF of the first

layer, out of 80, was calibrated relative to the hydration free energy of the permeant, whereas the PMF of the remaining 79 layers was calibrated to set the minimum Δ*G* to 0. The permeation resistance, *R*, through the 79 layers was obtained by multiplying *R* through one layer by a factor 79. The permeation resistances from the two steps were summed in order to obtain the permeability coefficient through the system. The resulting permeability coefficient calculation was:

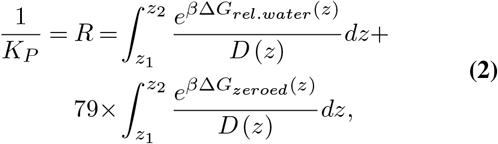

where Δ*G*_*rel*.*water*_ is the free energy relative to the hydration free energy and Δ*G*_*zeroed*_ is the free energy shifted so that the PMF minimum is 0.

The AWH simulation estimates a friction metric tensor *g*(*z*) (49), which can be used to derive the diffusion coefficient across the barrier structure, *D*(*z*) = *g*^−1^(z). To reduce the noise of the local diffusion coefficient curves, an 0.2 nm wide rolling median filter was applied. When symmetrizing the sampling, along the spatial dimension, by using the absolute coordinate values and accounting for the AWH bias across the sampling boundaries there are usually artificial spikes at the edges of the PMFs. These were been removed by setting the two lowest (0.005 nm and 0.015 nm) and highest (5.200 nm and 5.210 nm) points in the PMF to the value of their neighbours (0.025 nm and 5.19 nm, respectively). These minor adjustments had no effects on the calculated permeability coefficients.

The free energy profile through the skin’s barrier structure was calculated using a two-dimensional AWH setup, using a harmonic potential to steer the permeant across the system, also referred to as the *Z* dimension, and an alchemical free energy reaction coordinate(8). This allows sampling the free energy along the permeation direction and also the relative insertion free energy of the permeant from vacuum. In turn, this enables a direct calibration to the hydration free energy, since the vacuum state is the same in both cases. Like when calculating the hydration free energy, the estimated AWH initial error was set to 10 kJ mol^−1^. The AWH input diffusion constant was set to 3 × 10^−5^ nm^2^ ps^−1^ for the spatial pulling dimension and 5 × 10^−5^ ps^−1^ along the alchemical free energy dimension. The input diffusion constant affects only the AWH histogram size, it does not affect the computed diffusion coefficient, which is obtained from the AWH friction metric during data analysis. The AWH force constant along the spatial *Z* dimension (normal to the lamellar stack), steering the permeant relative to the ceramide fatty acid chains using a harmonic pull potential, was set to 25000 kJ mol^−1^ nm^−2^. The force constant also determines the resolution along the reaction coordinate dimension. For each permeant, four to six (see Table S1) sets of simulations were run with heavy hydrogen atoms (see above) and a 3 fs integration time step. These were run using 24 communicating walkers, each running for 450 ns. The covering check in the initial AWH stage only took into account simulations that covered the whole alchemical dimension and at least a diameter of 0.8 nm along the spatial dimension. From these simulations a combined diffusion coefficient was calculated using the AWH friction metric from all contributing walkers. The combined PMF was derived from the average of the independent PMFs from the four to six sets of simulations. The combined outputs were used to calculate the permeability coefficients presented in Table S1.

Along the alchemical free energy dimension it is the end states, i.e., the fully interacting and fully decoupled states, that are of highest interest, as the difference in free energy between them corresponds to the probability of transferring the permeant from vacuum into the skin’s barrier structure. Therefore, the target distribution used in these simulations put more weight on the end states, especially the state with interactions fully turned on. The target distribution along the alchemical free energy dimension that was used is shown in Fig. S4. Along the spatial pulling dimension the target distribution was uniform.

One-dimensional AWH simulations of testosterone across the barrier structure were also performed, for evaluation and comparison to the two-dimensional AWH results. They were run using four independent sets of simulations, each using 24 communicating AWH walkers running for 600 ns. The simulation settings were the same as specified above, except that there was no alchemical free energy dimension.

## Results

### Permeability Coefficient Calculations

Running one-dimensional AWH simulations, with only a spatial reaction coordinate dimension across the barrier structure, did not sample the free energy profile of testosterone efficiently. None out of four sets of simulations, each consisting of 24 communicating walkers sharing the same AWH bias and running for 600 ns for a total simulation time of 14.4 μs in each of the four independent simulation sets, sampled enough to cover the reaction coordinate with the condition that a single walker must cover a diameter of 0.8 nm in order to be taken into account. It is possible that this requirement was too strict for a system with inherently slow diffusion. This meant that the one-dimensional AWH simulations did not leave the AWH histogram equilibration stage, as part of the AWH initial stage. Increasing the AWH input diffusion constant would have increased the bias more quickly, meaning covering of the reaction coordinate would have been quicker. However, that might also have resulted in too high free energy barriers in the PMF. The average PMF from the four sets of 1D AWH simulations is compared to the results from the 2D AWH simulations, including an alchemical dimension, in Fig. 3. The results from 58 μs of 1D AWH were worse, assuming that the 55 μs of 2D AWH simulations are closest to convergence, than after 11 μs of 2D AWH with an alchemical reaction coordinate, and fairly similar to the FR pulling simulations shown in Fig. S3, but with higher uncertainty, i.e. a larger difference between the four sets of simulations.

**Fig. 3.**
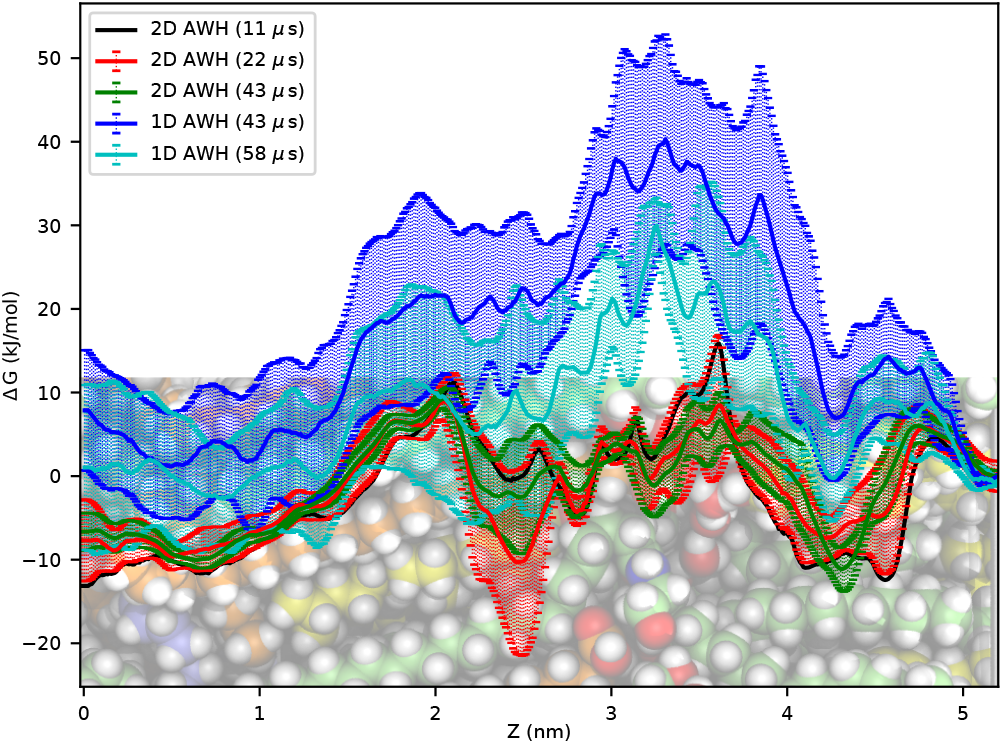
PMFs of testosterone comparing the 2D spatial/alchemical AWH (in the fully interacting alchemical *λ* state) results to 1D AWH with only a spatial reaction coordinate dimension. In both cases the PMFs are symmetrized and only half the PMFs are presented and the PMFs are calibrated to 0 at the ceramide sphingoid chain interface (5.2 nm), to make comparisons easier. The uncertainties represent one standard error of the mean. The 11 μs 2D AWH results are from one set of 24 communicating walkers. AWH analyses do not give a reliable error estimate from one set of simulations, therefore there is no error presented for the 11 μs AWH plot (in black). The 1D AWH simulations of 58 μs required approximately the same computation time as 22 μs of 2D AWH. A snapshot of the molecular system is shown at the lower portion of the plot to indicate where the head groups are located (at ≈3.2 nm to 3.3 nm).

The results from the two-dimensional AWH simulations agreed well with the experimental data, thanks to the improved sampling of the free energy landscape. The relationship between calculated and experimental permeability coefficients are presented in Fig. 4. The numerical results are presented in Table S1. The largest outlier was urea, followed by hydromorphone with log K_P_ deviations of − 2.4 and 1.5 cmh^−1^, respectively. The average absolute log K_P_ deviation was 0.8 cmh^−1^ (see Table S1). If excluding urea and hydromorphone it would be as low as 0.4 cmh^−1^.

**Fig. 4.**
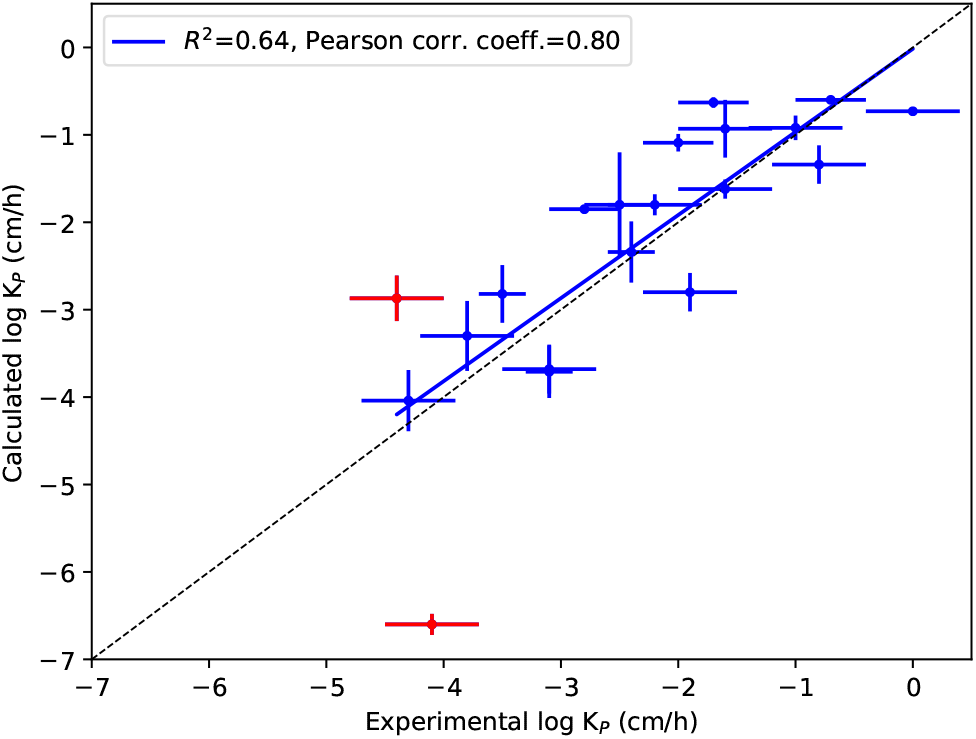
The correlation between experimental log K_P_ and those calculated from MD simulations. The blue line shows the linear regression, whereas the dashed black line is the identity line. The two largest outliers, urea and hydromorphone, are marked in red (still included in the linear regression). The error bars represent 1 SEM (standard error of the mean), which is approximated for experimental values. Details about the approximation is presented in the header of Table S1.

In the supplementary material (Figs. S5 and S6) the PMFs and local diffusion coefficients across the skin’s barrier structure are shown. There are also comparisons to our previously published results (7) using FR pulling.

#### Convergence of Diffusion Coefficients and PMFs

We have observed that the calculated diffusion coefficient depends on the simulation time, with lower diffusion coefficient the longer the simulation time. In general, the same trend is also observed regarding the PMF, i.e., lower free energy barriers with more sampling. We have seen this using all MD simulation sampling methods mentioned herein: AWH (1D with a spatial dimension and 2D with an added alchemical free energy perturbation dimension), umbrella sampling and FR pulling. Most simulations in this study were run for 450 ns (with 24 communicating AWH walkers). Our observations are that the calculated permeability coefficients are largely unaffected when increasing the simulation time by a factor two, from 450 ns to 900 ns per AWH walker (see Table S2 and Fig. S9).

## Discussion

When comparing PMFs of testosterone from the two-dimensional AWH simulations performed in this study to previously performed (unpublished) FR pulling simulations (Fig. S3), it is clear that the PMFs from the 30 μs pulling simulations are closer to the PMFs from AWH. The calculated log K_P_ for testosterone from the FR pulling simulations (taking the 80 stacked lipid bilayers with the same two-step process as described above when calculating the permeability coefficient) was −4.6 cmh^−1^, compared to −1.8 cmh^−1^ for the AWH simulations and − 2.5 cmh^−1^ for the *ex vivo/in vitro* measurements (with reports ranging from − 1.9 cmh^−1^ to 3.2 cmh^−1^(50–53), corrected to a temperature of 305 K).

The two-dimensional AWH sampling, with one alchemical dimension, was also more efficient than using a one-dimensional approach with only a spatial reaction coordinate, as shown in Fig. 3. The main drawback with an alchemical free energy reaction coordinate dimension is that the simulation throughput is lower compared to FR pulling simulations as well as one-dimensional AWH simulations using only a spatial coordinate. However, the possibility to more accurately sample the free energy landscape makes it attractive for problems that are difficult to address with conventional methods, such as permeability coefficient predictions in gel-phase systems, like the skin’s barrier structure.

Another advantage is that it is easier to prepare permeability calculation simulations using an alchemical dimension than with conventional procedures, as the permeant molecule can be inserted anywhere in the system with all interactions turned off. The AWH simulations will then turn on the interactions where favourable. On the contrary, with standard FR pulling, as well as with one-dimensional AWH pulling or umbrella simulations, the molecule must be grown into the system and/or pulled to suitable starting positions, followed by equilibration of the simulation system, before the production simulations are started. To determine the free energy landscape without an alchemical dimension in the calculation, it would also be necessary to know the free energy of inserting the molecular for at least one position along the barrier structure; otherwise the PMF cannot be calibrated compared to a solution or formulation.

The permeability coefficient calculations presented herein only account for the passage of a chemical compound across the skin’s barrier structure. Lateral diffusion within the skin’s barrier structure is not calculated, and permeation through the stratum corneum cell interiors, as well as through viable cell layers of epidermis, are ignored. While these are admittedly approximations, the contributions from these factors are expected to be negligible in comparison to the permeation resistance across the skin’s barrier structure for all but very lipophilic compounds.(54) Second, it is foremostly the permeability across the skin’s barrier structure that can be tuned using chemical permeation enhancers, and thereby exploited in transdermal drug delivery design. The two-step process to calculate the permeability coefficient, with partitioning from the delivery vehicle into the lipid barrier followed by permeation through the remaining layers of the lipid barrier, is a simplification of the process, but has given good agreement with experimental measurements. In practice, it only affects the calculated permeability of hydrophilic permeants, when using water as the delivery vehicle, compared to setting the PMF minimum to 0 for all permeation barrier layers.

As can be inferred from Wang and Klauda (22) and from Scheuplein and Blank (54) as well as from Eq. 20 in Marrink and Berendsen (48), and discussed in the Introduction section above, the permeability coefficient does not correspond to an average permeation speed through the skin, but will become lower the thicker the skin sample. Therefore, the permeability coefficient through the skin’s barrier structure alone would be higher than through full thickness skin samples used in *ex vivo/in vitro* permeability measurements. We would therefore expect calculated permeability coefficients, referring exclusively to the skin’s barrier structure, to be slightly higher than those measured *ex vivo/in vitro* through full thickness skin, assuming that the skin samples are intact. Our calculated log K_P_ is 0.1 cmh^−1^ higher than experimental results on average, and 0.2 cmh^−1^ higher if excluding urea and hydromorphone — the largest outliers.

Measuring skin permeability using diffusion cells, such as Franz cells, require long experiment times. How long depends on the permeant’s lag time.(55) It is common to run experiments for 24 h to 72 h.(55) In extreme cases, with low permeability and long lag times, this might even be insufficient. However, long-time exposure to water leads to skin degradation and increased skin permeability.(55, 56) These factors can make it difficult to compare theoretical results with those obtained *ex vivo/in vitro*, since the *ex vivo/in vitro* measurements may sometimes be too short for the permeation lag time and too long for keeping the skin sample intact. The largest outliers in this study (see Table S1) are urea (log K_P_ diff. −2.5 cmh^−1^) and hydromorphone (log K_P_ diff. 1.5 cmh^−1^), followed by salicylic acid and ethanol. The correlation plot of the results, excluding urea and hydromorphone, is shown in Fig. S7. The permeation enhancing effects of urea (57) were not taken into account in these simulations, which could be a reason for the discrepancy. For hydromorphone, there is a remarkable similarity between the average of three different predicted log K_P_ obtained using mathematical QSPR models of −3.2 cmh^−1^ (10, 58–60) and our predicted log K_P_ at 32 °C of −2.9 cmh^−1^, contrasting with the experimentally obtained value of −4.8 cmh^−1^ at 37 °C (61), with no corrections for pH/pK_a_. This discrepancy between predicted and experimental data may call for reinvestigation of the experimentally measured permeability coefficient of hydromorphone, preferably from multiple laboratories, although it could of course also indicate permeation properties not picked up by QSPR models either.

## Conclusion

We have shown that the model of the skin’s barrier structure that was proposed a few years ago (6) can be used to accurately predict permeability coefficients of a large spectrum of molecules, with clearly better computational efficiency than previous methods.

To our knowledge, this is the first report describing the use of a two-dimensional reaction coordinate with one spatial dimension and one alchemical free energy dimension. It is also the first study to employ the Accelerated Weight Histogram method to sample such a free energy landscape. The obtained PMFs are detailed and some of their features only hinted at when using forward-reverse pulling simulations with very slow pulling speeds. When taking the multiple bilayers of the skin’s barrier structure into account, our previously published calculations underestimated the skin permeability coefficients (7), largely explained by the inability of the forward-reverse pulling method to properly sample the free energy landscape.

Experimental permeability coefficient measurements are associated with variations between laboratories as well as with inter- and intraindividual differences. This makes it treacherous to strictly compare calculated permeability coefficients with experimental data. This needs, particularly, to be taken into account when creating QSPR models.(4) A suggested approach for building a reliable QSPR model would therefore be that either all training data should come from one laboratory using the same experimental setup for all compounds or, preferably, there should be measurements from several laboratories, using the same methods, for each compound. When predicting permeability coefficients using MD simulation there is no need for training the method with experimental data, but reliable experimental measurements are required for verifying the techniques and simulation results. The correlation between calculated and experimental permeability coefficients presented here proves the usefulness of MD simulation for transdermal drug delivery design, although the method’s precision may presently not always allow for differentiation of very similar molecules.

Perhaps the most important for transdermal drug delivery design is that MD simulation can explain experimental *ex vivo/in vitro* data by predicting where in the skin’s barrier structure the main permeation barriers are located for each specific permeant. This knowledge, along with the PMFs (in effect partition profiles) of chemical permeation enhancers, can help selecting suitable drug delivery formulation excipient combinations tailored for each specific API.

## ACKNOWLEDGEMENTS

The computations were enabled by resources provided by the Swedish National Infrastructure for Computing (SNIC 2020/5-220 and 2021/5-217) at PDC Centre for High Performance Computing at KTH and NSC at Linköping University as well as internal resources from ERCO Pharma AB and Erik Lindahl’s research group.

## DATA AVAILABILITY

The input data and parameters are available for download from https://doi.org/10.5281/zenodo.5883411.

## AUTHOR CONTRIBUTIONS

Lundborg, Norlén, Wennberg and Lindahl designed the research. Lundborg, Lidmar, Hess and Wennberg developed the simulation protocols. Lundborg carried out all simulations and analyzed the data with assistance from Wennberg. All authors wrote the article, with Lundborg being the primary author.

## COMPETING FINANCIAL INTERESTS

This work was funded by ERCO Pharma AB.

ERCO Pharma AB has submitted a patent application covering using this lipid barrier model for skin permeability calculations (application number PCT/EP2017/076237 and title “SKIN PERMEABILITY PREDICTION”).

Norlén, Lundborg and Wennberg have stock options in ERCO Pharma AB. Lundborg and Wennberg are employed by ERCO Pharma AB.

## Footnotes

1 Relative composition in: molar % ceramides/molar % cholesterol/molar % free fatty acids/relative amount of cholesterol on ceramide sphingoid side/molar % acyl ceramide EOS (included in the relative ceramide concentration)/water molecules per lipid (not included in the molar % concentrations of the lipids).

## Notes

https://doi.org/10.5281/zenodo.5883411

