## Supplementary Information for "Predictions of Skin Permeability Using Molecular Dynamics Simulation from Two-Dimensional Sampling of Spatial and Alchemical Perturbation Reaction Coordinates"

### Supplementary Note S1: Methods

When running AWH simulations it is possible to run multiple simulations in parallel, referred to as AWH walkers. The simulations can share their sampling, and thereby use the same bias, by communicating at regular intervals. This can improve the total simulation throughput compared to running a single simulation for a longer time. For the simulations of the lipid barrier structure 24 MD simulations with communicating AWH walkers were run, whereas 16 communicating AWH walkers were used for the hydration free energy calculations. The AWH histograms were equilibrated as part of the initial stage, using the criterion that at least 80 % of the target region was sampled within 20 % of the target distribution.

During the course of this project it was discovered that GROMACS 2021 running AWH with an alchemical free energy dimension did not properly take PME contributions into account along the alchemical dimension when calculating PME on a GPU accelerator. After this bug was fixed all hydration free energy calculations were rerun as well as one setup, with 24 communicating AWH walkers, each running 450 ns, calculating the permeability through the barrier. It was found that a systematic shift in the PMF, relative to the vacuum state, in the range of $0\,\mathrm{kJ\,mol^{-1}}$ to $13\,\mathrm{kJ\,mol^{-1}}$ depending on the permeant, was the only significant difference. The PMF results from the GROMACS version with the bug were shifted to align with the new correct output.

### Supplementary Note S2: Results

**S2.1. Comparisons to other MD simulation methods.** In the data presented in this section we pulled testosterone using a constraint, i.e., an infinitely stiff spring, to avoid having to select a suitable spring constant for each permeant (1). Compared to the 2018 study we also equilibrated the system longer after inserting the two copies of the permeant into the system, (20 ns instead of 1 ns). As can be seen in Fig. S1 the results, for testosterone, were very dependent on the pulling speeds. As these results are not the focus of this study — they are just included for comparison — we will refer to the original paper(2) for a more thorough description of the methods.

We also evaluated umbrella sampling simulations for calculating the PMFs. We inserted four copies of testosterone, evenly distributed along the reaction coordinate, and grew them in, by turning on their interactions with their environment, over a period of 1 ns. This was repeated in 80 steps along the reaction coordinate, giving 320 umbrella sampling positions at random lateral positions. During the umbrella simulations the permeants were held in place using a spring constant of $5000\,\mathrm{kJ\,mol^{-1}\,nm^{-2}}$. The 80 simulations were run for 150 ns (for a total of 12 000 ns). To estimate the equilibration times (how much to discard in the beginning of the simulations) a time window of 100 ns, with a shifting starting time, was analyzed. As can be seen in Fig. S2, even if the difference between the shortest and longest equilibration times was not statistically significant, there was an obvious systematic drift in the PMF with increasing equilibration times, most clearly seen in Fig. S2b. This indicated that discarding 50 ns (out of 150 ns) might not be enough. While, in the case shown here, the effect is not very large for the symmetrized PMF (Fig. S2a) the long times to reach convergence was not very attractive. Comparing Fig. S2a) to the previously published results from FR pulling simulations(2), shown by the red curve in the Testosterone sub figure in Fig. S5, indicates that the height of the free energy profile from these umbrella simulations roughly matched the ones from FR pulling at a pulling speed of $0.2\,\mathrm{nm\,ns^{-1}}$. However, they required approximately the same total simulation time as the pulling simulations at $0.005\,\mathrm{nm\,ns^{-1}}$. We expect that the umbrella sampling simulations can be further optimized, and do not claim that umbrella sampling is better or worse than FR pulling. Both methods face similar challenges with the long correlation times of the skin's barrier system existing in

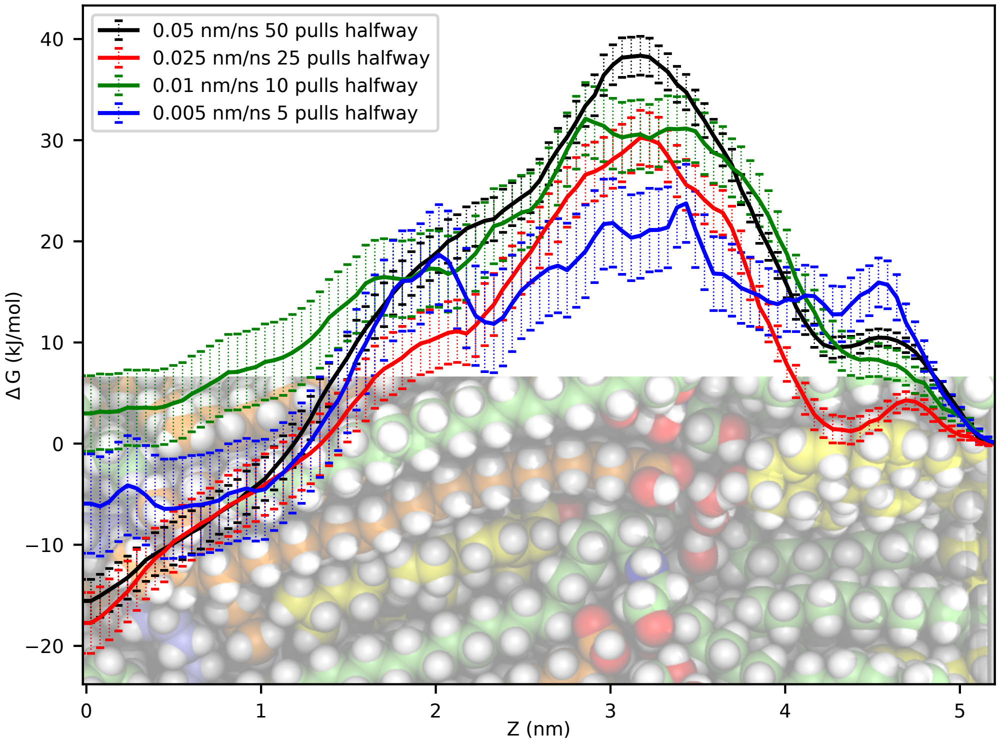


**Fig. S1.** PMFs of testosterone, from forward-reverse pulling simulations, with different pulling speeds. The results are from $10\,\mu s$ of sampling and the PMFs are calibrated to 0 at the ceramide sphingoid chain interface ($5.2\,nm$), to make comparisons easier. The general trend is that slower pulling speeds give lower free energy barriers and more detailed PMFs. However, fewer pulls also means a larger variability. Two permeants were pulled halfway through the system in order to shorten the simulations. A snapshot of the molecular system is shown at the lower portion of the plot to indicate where the head groups are located (at $\approx 3.2\,nm$ to $3.3\,nm$).

a gel-like state.

**S2.2. Comparison of testosterone PMFs from 2D AWH to FR Pulling.** Using the slowest pulling speed shown in Fig. S1 we compared the data from FR pulling simulations at $10$ and $30\,\mu s$ to the two-dimensional AWH setup, that was used for this project, at $11$ , $22$ and $43\,\mu s$. The results are presented in Fig. S3 and show that the differences were small between the AWH simulations at $22$ and $43\,\mu s$ and that the results from $30\,\mu s$ FR pulling were further away from the final AWH results than the results after $11\,\mu s$ of AWH simulations. It can also be noted that the results from FR pulling simulations seem to approach those from AWH, but since the output from $30\,\mu s$ of FR pulling is still far from the AWH results it can be concluded that many more FR pulling simulations, in order to achieve a longer total simulation time, would be needed to come close to the AWH results. Looking also at the testosterone diffusion coefficient profile in Fig. S6 it can be concluded that the largest differences between FR pulling and AWH are observed in the region with the slowest diffusion, i.e., $Z \sim 2.5\,nm$ to $\sim 4.5\,nm$.

**S2.3. Permeability Coefficient Calculations.** The permeability coefficients were calculated using Eq. 1 (in the main paper) using the PMF and diffusion coefficient profiles presented in Figs. S5 and S6. The results are presented in Table S1 and in Fig. 4 (in the main paper).

In this study we have been focusing on the permeability coefficients of neutral (non-ionized) forms of the permeants. There were eight molecules that are ionizable under the *ex vivo/in vitro* conditions: codeine, diclofenac, hydromorphone, ibuprofen, lidocaine, naproxen, nicotine and salicylic acid. For most of them there are published permeability coefficients available, whereas for codeine and hydromorphone the corrections by Abraham and Martins (4) were used. In order to examine whether the neutral forms of ionizable permeants were in agreement with experimental results we divided the molecules into two groups, separating those that are not ionizable under the *ex vivo/in vitro* conditions from those that are ionizable. The correlation plots from these analyses are presented in Fig. S8 and show that there is no clear difference between the two groups, but the permeability coefficients of the set of non-ionizable compounds are better predicted by MD simulations.

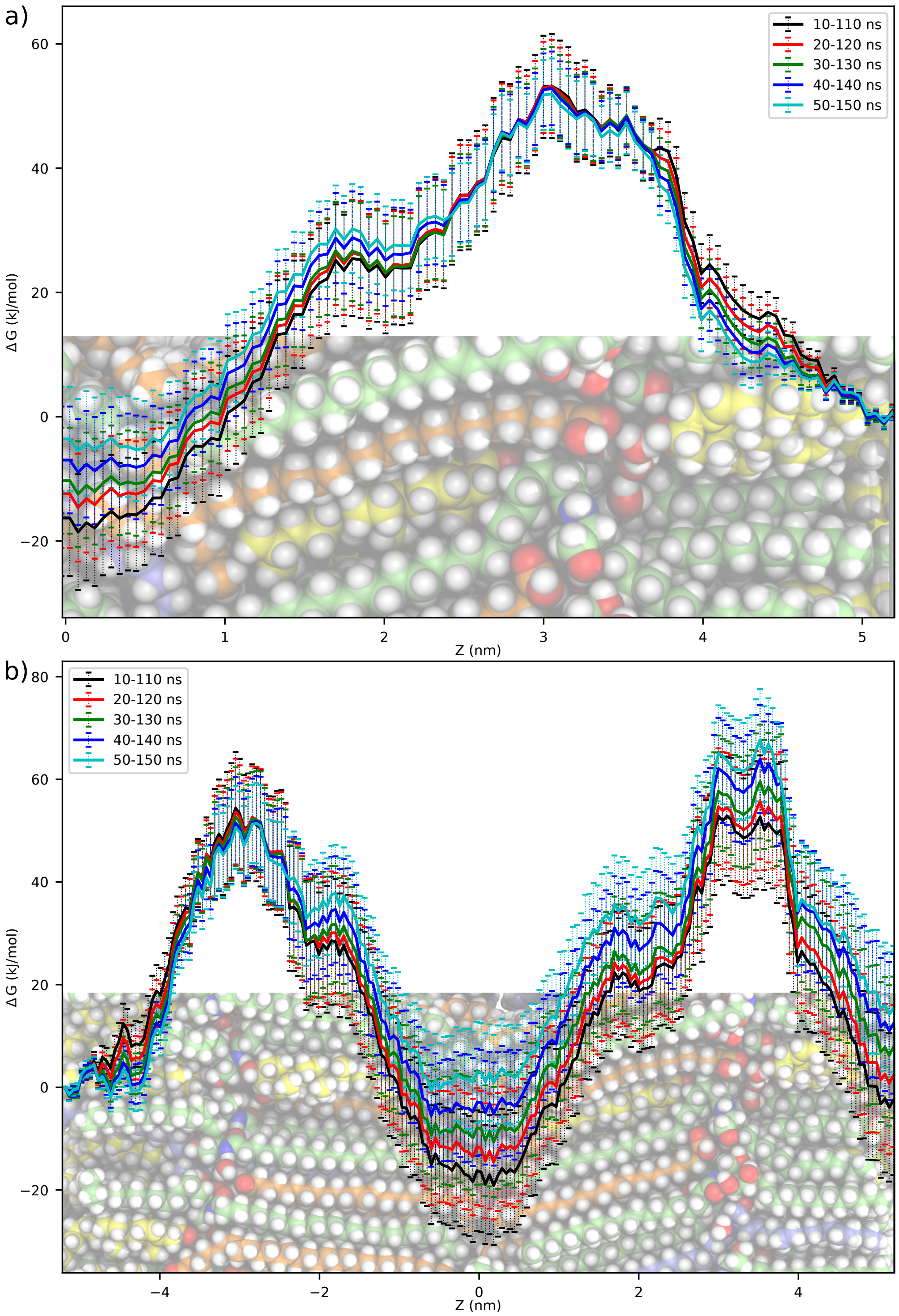


**Fig. S2.** PMFs of testosterone from umbrella sampling simulations with increasing equilibration time. In a) the data is enforced to be periodic, with connected end points, and symmetrized. The PMFs are calibrated to 0 at the ceramide sphingoid chain interface ($5.2\,\mathrm{nm}$), to make comparisons easier. In b) the data is not symmetrized without connected end points (the PMF is set to 0 only at $-5.2\,\mathrm{nm}$). A snapshot of the molecular system is shown at the lower portions of the plots to indicate where the head groups are located (at $\approx\pm3.2\,\mathrm{nm}$ to $\pm3.3\,\mathrm{nm}$).

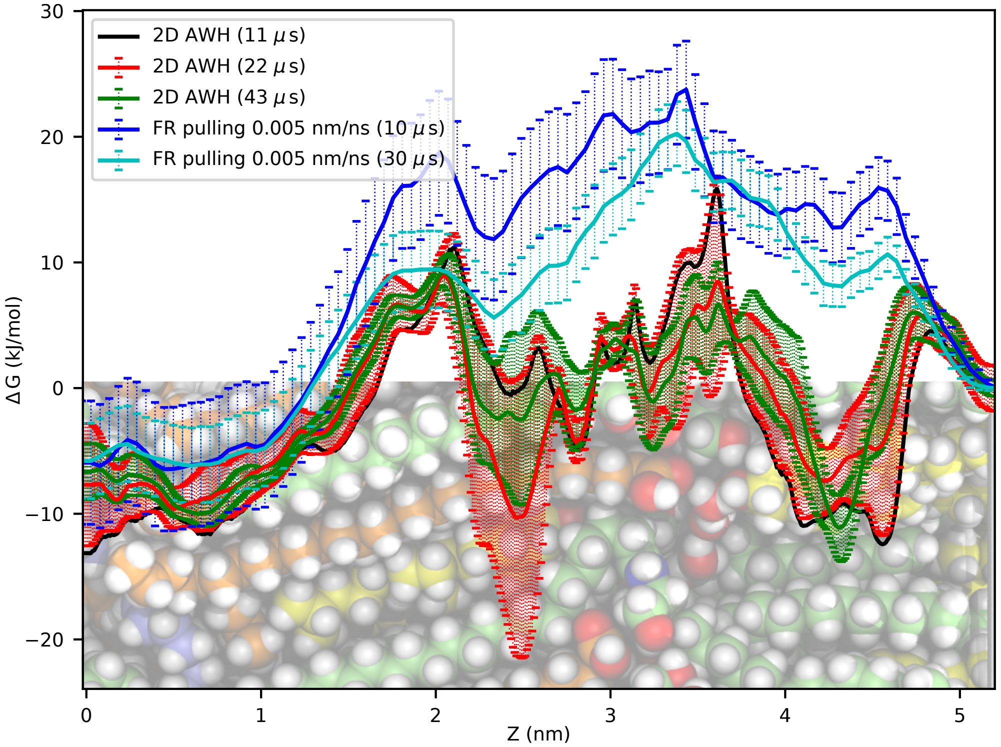


**Fig. S3.** PMFs of testosterone comparing the 2D spatial/alchemical AWH (in the fully interacting alchemical $\lambda$ state) results to slow forward-reverse pulling experiments. In both cases the PMFs are symmetrized and only half the PMFs are presented and the PMFs are calibrated to 0 at the ceramide sphingoid chain interface ($5.2\,\mathrm{nm}$), to make comparisons easier. The $11\,\mu\mathrm{s}$ AWH results are from one set of 24 communicating walkers. AWH analyses do not give a reliable error estimate from one set of simulations, therefore there is no error presented for the $11\,\mu\mathrm{s}$ AWH plot (in black). In the pulling experiments two permeants were pulled halfways through the system in order to shorten the simulations. FR pulling simulations of $30\,\mu\mathrm{s}$ required approximately the same computation time as $11\,\mu\mathrm{s}$ of 2D AWH. A snapshot of the molecular system is shown at the lower portion of the plot to indicate where the head groups are located (at $\approx 3.2\,\mathrm{nm}$ to $3.3\,\mathrm{nm}$).

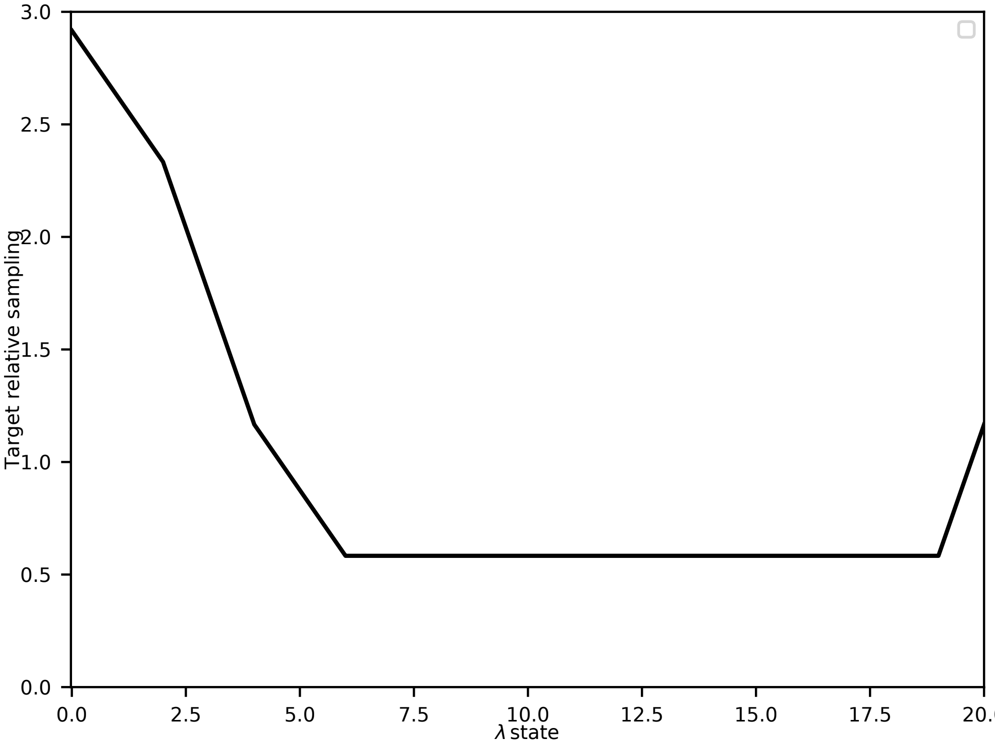


**Fig. S4.** AWH target distribution along the alchemical free energy axis. $\lambda$ state 0 was fully interacting and at $\lambda$ state 20 there were no interactions between the molecule and its surroundings. Along the pull axis the target distribution was uniform.

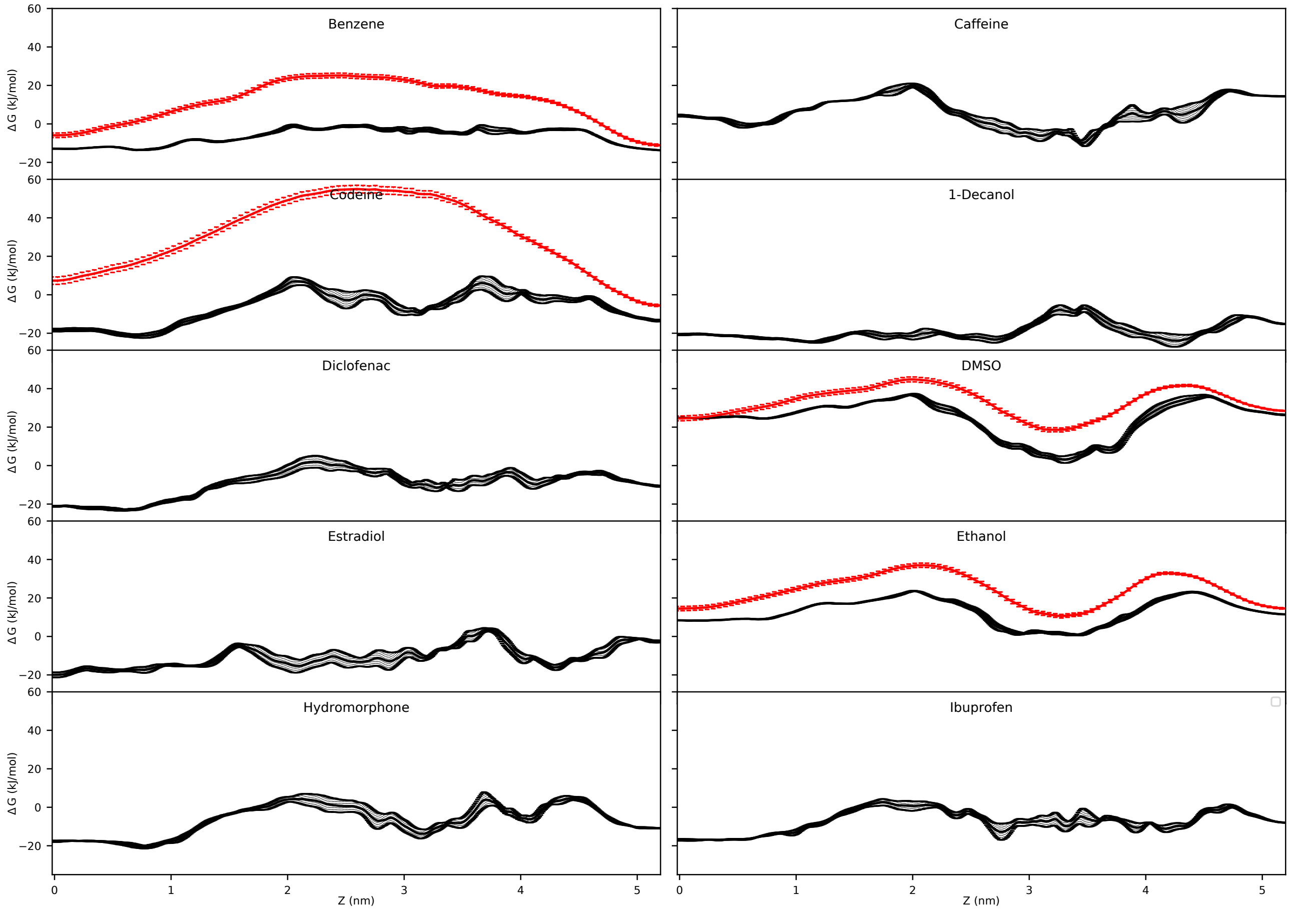


**Fig. S5.** PMFs of 20 permeants. The PMFs are calibrated related to the hydration free energies of the compounds. The profiles in black show the results from this study, whereas the profiles in red are from our previous FR pulling simulations (pulling at $0.2\,\mathrm{nm\,ns^{-1}}$) (2, 3). The curves in red were only calibrated based on the free energy difference, to the water solvent, at the ceramide sphingoid chain interface region (at $\sim 5.2\,\mathrm{nm}$). *Continued on next page.*

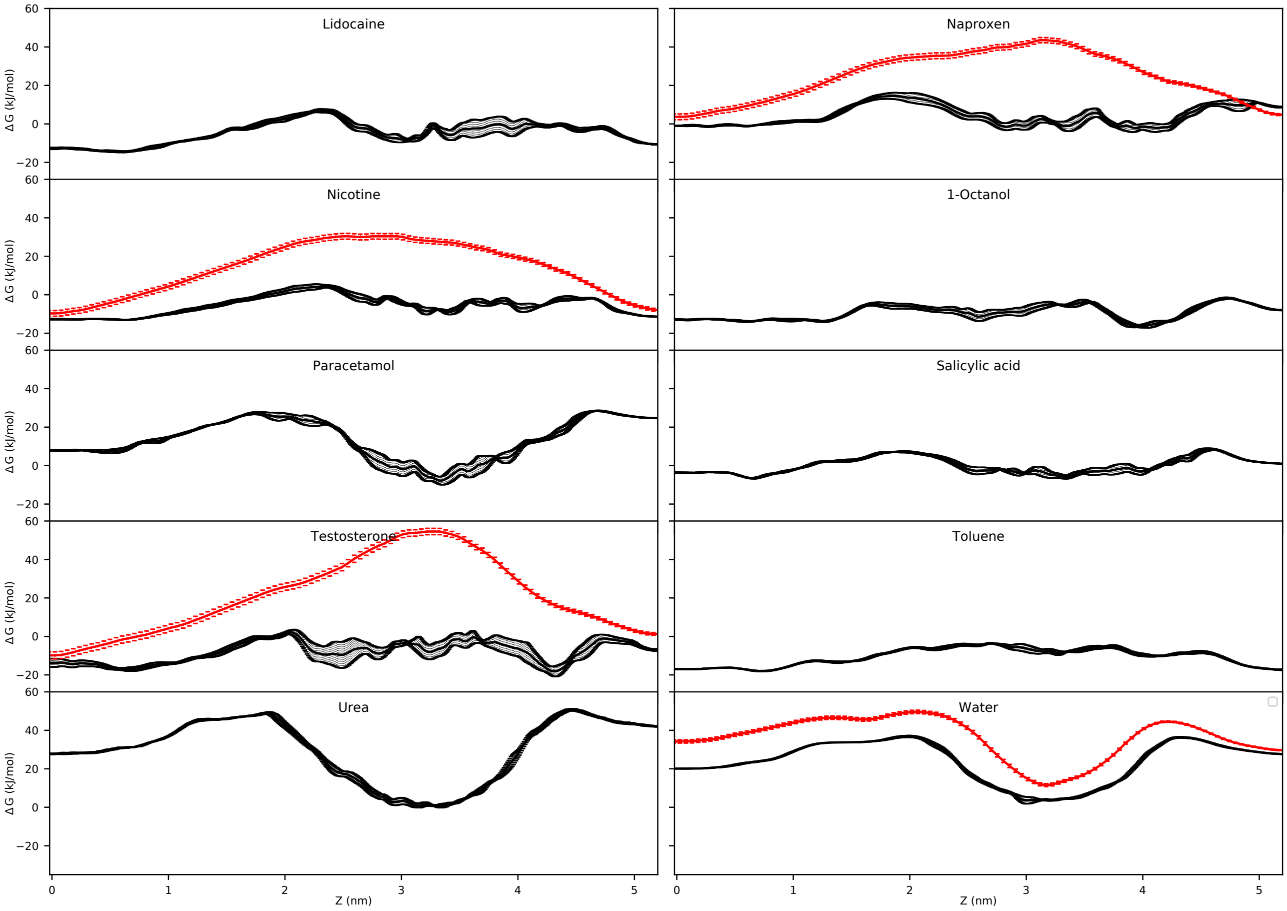


**Fig. S5.** PMFs of 20 permeants. The PMFs are calibrated related to the hydration free energies of the compounds. The profiles in black show the results from this study, whereas the profiles in red are from our previous FR pulling simulations (pulling at $0.2\,\mathrm{nm\,ns^{-1}}$) (2, 3). The curves in red were only calibrated based on the free energy difference, to the water solvent, at the ceramide sphingoid chain interface region (at $\sim 5.2\,\mathrm{nm}$). *Continued from previous page.*

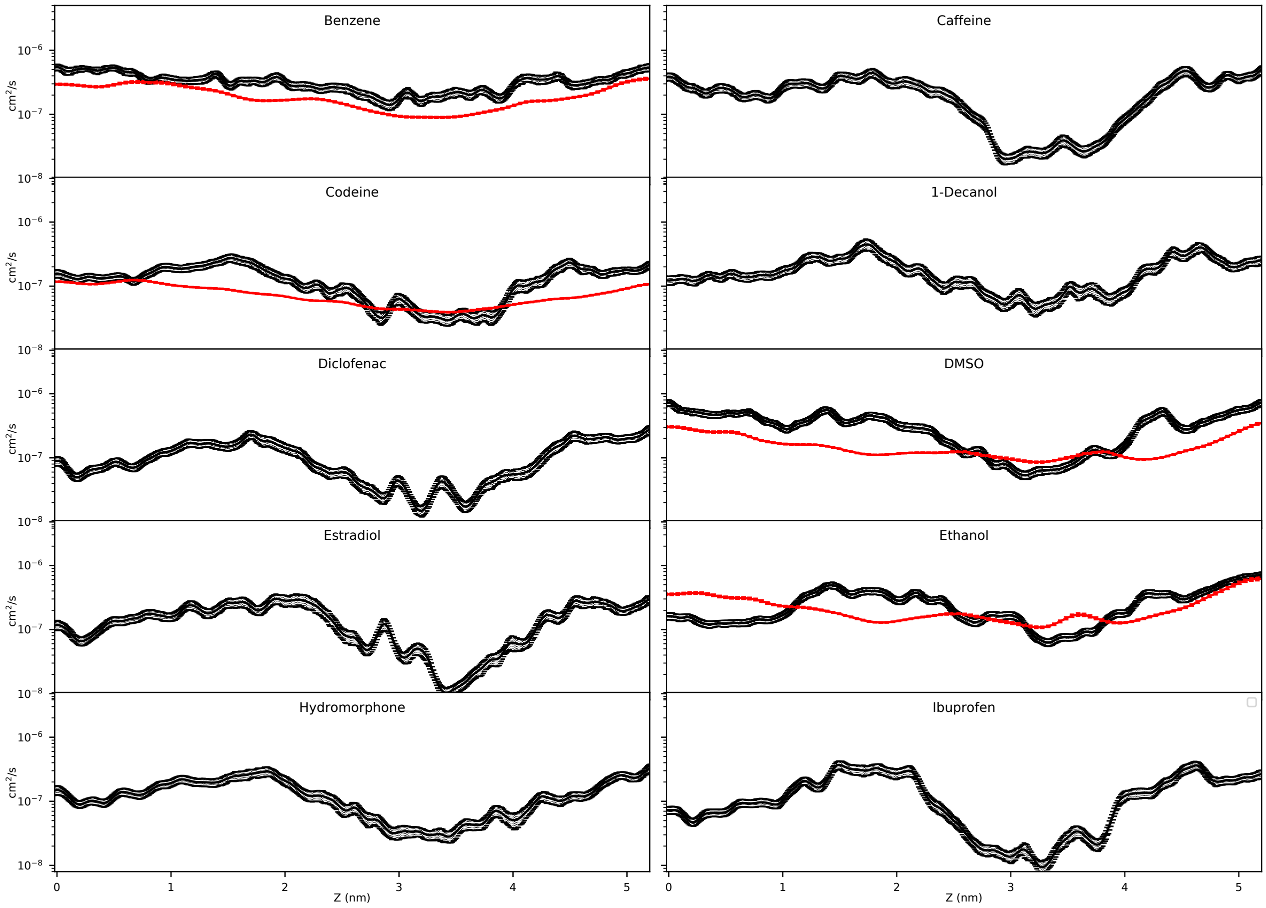


**Fig. S6.** Diffusion coefficients of 20 permeants. The profiles in black show the results from this study, whereas the profiles in red are from our previous FR pulling simulations (pulling at $0.2\,\mathrm{nm\,ns^{-1}}$) (2, 3). The diffusion profiles from AWH were smoothened using a running median filter of $0.2\,\mathrm{nm}$ width, whereas a Hanning filter of $11\,\mathrm{point}$ ($0.5\,\mathrm{nm}$) width was used for the diffusion profiles from FR pulling. *Continued on next page.*

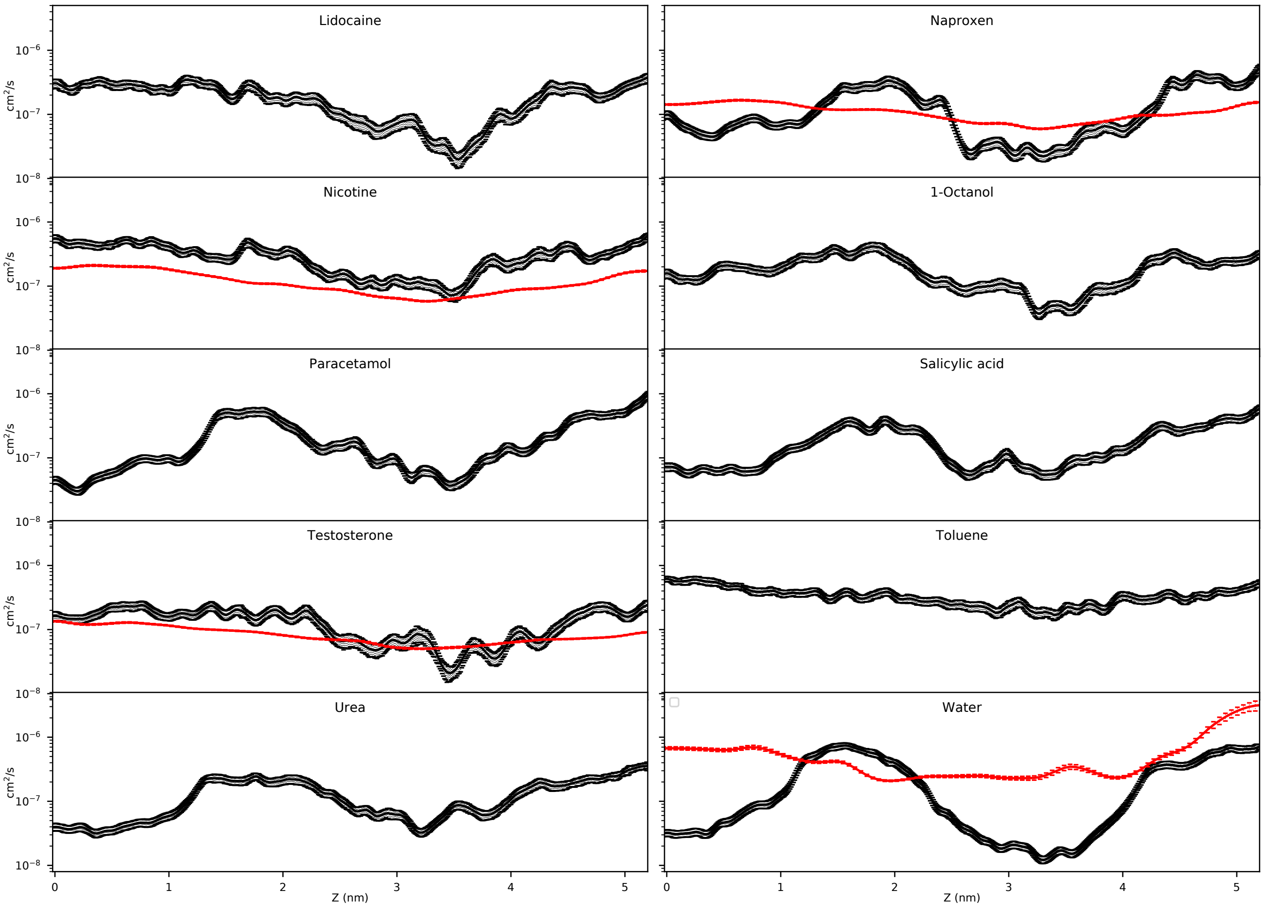


**Fig. S6.** Diffusion coefficients of 20 permeants. The profiles in black show the results from this study, whereas the profiles in red are from our previous FR pulling simulations (pulling at $0.2\,\mathrm{nm\,ns^{-1}}$) (2, 3). The diffusion profiles from AWH were smoothened using a running median filter of $0.2\,\mathrm{nm}$ width, whereas a Hanning filter of $11\,\mathrm{point}$ ($0.5\,\mathrm{nm}$) width was used for the diffusion profiles from FR pulling. *Continued from previous page.*

**Table S1.** Permeability coefficients of unionized species at 32 ℃, with corrections for temperature differences according to Abraham and Martins (4). If only observations from ionized species were found, corrections for pH/p$K_a$ were then obtained from Abraham and Martins (4). The log $K_{P_{exp}}$ uncertainty is the standard error of the mean (SEM) of the measurements, if there were at least three reported measurements available. If there were only two experimental measurements the SEM was assumed to be 0.3 and shown in italics and if there was only one experimental measurement the SEM was reported as 0.4 (in italics). The column $N_{sim.}$ shows the number of independent sets of simulations used to calculate the permeability coefficient and its uncertainty.

| **Permeant** | **log $K_{P_{exp}}$** (cm/h) | **log $K_{P_{calc}}$** (cm/h) | **$N_{sim.}$** | **Tot. sim. time** ($\mu$s) |
|---|---|---|---|---|
| Benzene | $-0.7 \pm$ *0.3*[a] | $-0.6 \pm 0.0$ | 4 | 43.2 |
| Caffeine | $-3.5 \pm 0.2$[b] | $-2.8 \pm 0.3$ | 4 | 43.2 |
| Codeine | $-3.8 \pm$ *0.4*[c] | $-3.3 \pm 0.4$ | 6 | 64.8 |
| 1-Decanol | $-0.8 \pm$ *0.4*[d] | $-1.3 \pm 0.2$ | 4 | 43.2 |
| Diclofenac | $-1.9 \pm$ *0.4*[e] | $-2.8 \pm 0.4$ | 5 | 54.0 |
| DMSO | $-3.1 \pm$ *0.4*[f] | $-3.7 \pm 0.3$ | 5 | 54.0 |
| Estradiol | $-2.5 \pm 0.2$[g] | $-2.3 \pm 0.4$ | 5 | 54.0 |
| Ethanol | $-2.8 \pm$ *0.3*[h] | $-1.9 \pm 0.1$ | 5 | 54.0 |
| Hydromorphone | $-4.4 \pm$ *0.4*[i] | $-2.9 \pm 0.3$ | 5 | 54.0 |
| Ibuprofen | $-1.6 \pm$ *0.4*[j] | $-1.6 \pm 0.1$ | 4 | 43.2 |
| Lidocaine | $-2.2 \pm$ *0.4*[k] | $-1.8 \pm 0.1$ | 4 | 43.2 |
| Naproxen | $-1.6 \pm$ *0.4*[l] | $-0.9 \pm 0.3$ | 4 | 43.2 |
| Nicotine | $-2.0 \pm$ *0.3*[m] | $-1.1 \pm 0.1$ | 4 | 43.2 |
| 1-Octanol | $-1.0 \pm$ *0.4*[n] | $-0.9 \pm 0.1$ | 4 | 43.2 |
| Paracetamol | $-4.3 \pm$ *0.4*[o] | $-4.0 \pm 0.4$ | 4 | 43.2 |
| Salicylic acid | $-1.7 \pm$ *0.3*[p] | $-0.6 \pm 0.1$ | 4 | 43.2 |
| Testosterone | $-2.5 \pm 0.3$[q] | $-1.8 \pm 0.6$ | 4 | 43.2 |
| Toluene | $0.0 \pm$ *0.4*[r] | $-0.7 \pm 0.1$ | 4 | 43.2 |
| Urea | $-4.1 \pm$ *0.3*[s] | $-6.6 \pm 0.1$ | 4 | 43.2 |
| Water | $-3.1 \pm 0.2$[t] | $-3.7 \pm 0.3$ | 4 | 43.2 |
| Mean diff. | | 0.11 | | |
| Mean abs. diff. | | 0.69 | | |
| Mean sq. diff. | | 0.79[u] | | |

[a] log $K_{P_{exp}}$ $-0.80\,\mathrm{cm\,h^{-1}}$ at 25 °C (5) and $-0.95\,\mathrm{cm\,h^{-1}}$ at 31 °C (6).

[b] log $K_{P_{exp}}$ $-4.00\,\mathrm{cm\,h^{-1}}$ at 25 °C (5), $-3.59\,\mathrm{cm\,h^{-1}}$ at 32 °C (7) and $-3.19\,\mathrm{cm\,h^{-1}}$ at 32 °C (8).

[c] log $K_{P_{exp}}$ $-4.31\,\mathrm{cm\,h^{-1}}$ at 37 °C and pH 7.4 (9) (p$K_a$ 8.2).

[d] log $K_{P_{exp}}$ $-1.10\,\mathrm{cm\,h^{-1}}$ at 25 °C (10).

[e] log $K_{P_{exp}}$ $-1.74\,\mathrm{cm\,h^{-1}}$ at 37 °C, unionized (11).

[f] log $K_{P_{exp}}$ $-3.10\,\mathrm{cm\,h^{-1}}$ at 5 % conc. at unspecified temperature (12). Cf., at 40 % conc. the log $K_{P_{exp}}$ was $-1.80\,\mathrm{cm\,h^{-1}}$.

[g] log $K_{P_{exp}}$ $-2.01\,\mathrm{cm\,h^{-1}}$ at 37 °C (13), $-2.28\,\mathrm{cm\,h^{-1}}$ at 30 °C (5), $-2.34\,\mathrm{cm\,h^{-1}}$ at 34 °C (SC temperature) (14), $-2.37\,\mathrm{cm\,h^{-1}}$ at 32 °C (15), $-2.41\,\mathrm{cm\,h^{-1}}$ through full skin at 30 °C (16), $-2.49\,\mathrm{cm\,h^{-1}}$ at 25 °C (5) and $-3.52\,\mathrm{cm\,h^{-1}}$ at 26 °C (17).

[h] log $K_{P_{exp}}$ $-3.14\,\mathrm{cm\,h^{-1}}$ at 25 °C (10) and $-3.00\,\mathrm{cm\,h^{-1}}$ at 25 °C (18).

[i] log $K_{P_{exp}}$ $-4.82\,\mathrm{cm\,h^{-1}}$ at 37 °C, pH 7.4 (9) (p$K_a$ 8.2).

[j] log $K_{P_{exp}}$ $-1.44\,\mathrm{cm\,h^{-1}}$ at 37 °C, presumably unionized (19).

[k] log $K_{P_{exp}}$ $-2.15\,\mathrm{cm\,h^{-1}}$ at 32 °C, pH 10.0 (9) (p$K_a$ 8.0).

[l] log $K_{P_{exp}}$ $-1.42\,\mathrm{cm\,h^{-1}}$ at 37 °C, unionized (11).

[m] log $K_{P_{exp}}$ $-1.71\,\mathrm{cm\,h^{-1}}$ at unknown temperature, unionized (20), $-2.08\,\mathrm{cm\,h^{-1}}$ at 37 °C, unionized (21) .

[n] log $K_{P_{exp}}$ $-1.28\,\mathrm{cm\,h^{-1}}$ at 25 °C (10).

[o] log $K_{P_{exp}}$ $-4.30\,\mathrm{cm\,h^{-1}}$ at 32 °C (22).

[p] log $K_{P_{exp}}$ $-1.52\,\mathrm{cm\,h^{-1}}$ at 37 °C, unionized (11) and $-1.58\,\mathrm{cm\,h^{-1}}$ at 32 °C, pH 2.5 (23) (p$K_a$ 3.0).

[q] log $K_{P_{exp}}$ $-2.66\,\mathrm{cm\,h^{-1}}$ at 25 °C (5), $-3.40\,\mathrm{cm\,h^{-1}}$ at 26 °C (17), $-1.85\,\mathrm{cm\,h^{-1}}$ at 32 °C (24) and $-3.07\,\mathrm{cm\,h^{-1}}$ at 25 °C (25).

[r] log $K_{P_{exp}}$ $0.0\,\mathrm{cm\,h^{-1}}$ at unknown temperature (26).

[s] log $K_{P_{exp}}$ $-3.89\,\mathrm{cm\,h^{-1}}$ at 37 °C (27) and $-4.04\,\mathrm{cm\,h^{-1}}$ at 34 °C (SC temperature) (14).

[t] log $K_{P_{exp}}$ $-3.30\,\mathrm{cm\,h^{-1}}$ at 25 °C (10), $-3.70\,\mathrm{cm\,h^{-1}}$ at 25 °C (10), $-3.00\,\mathrm{cm\,h^{-1}}$ at 25 °C (18) and $-3.40\,\mathrm{cm\,h^{-1}}$ at unspecified temperature (12).

[u] $\mathrm{cm^2\,h^{-2}}$

**S2.4. Convergence of Diffusion Coefficients.** Fig. S9 shows how the calculated diffusion coefficients change with extended simulation times. Especially for testosterone it can be seen that longer simulation times result in lower diffusion coefficients. Table S2 shows the calculated permeability coefficients from the same simulations.

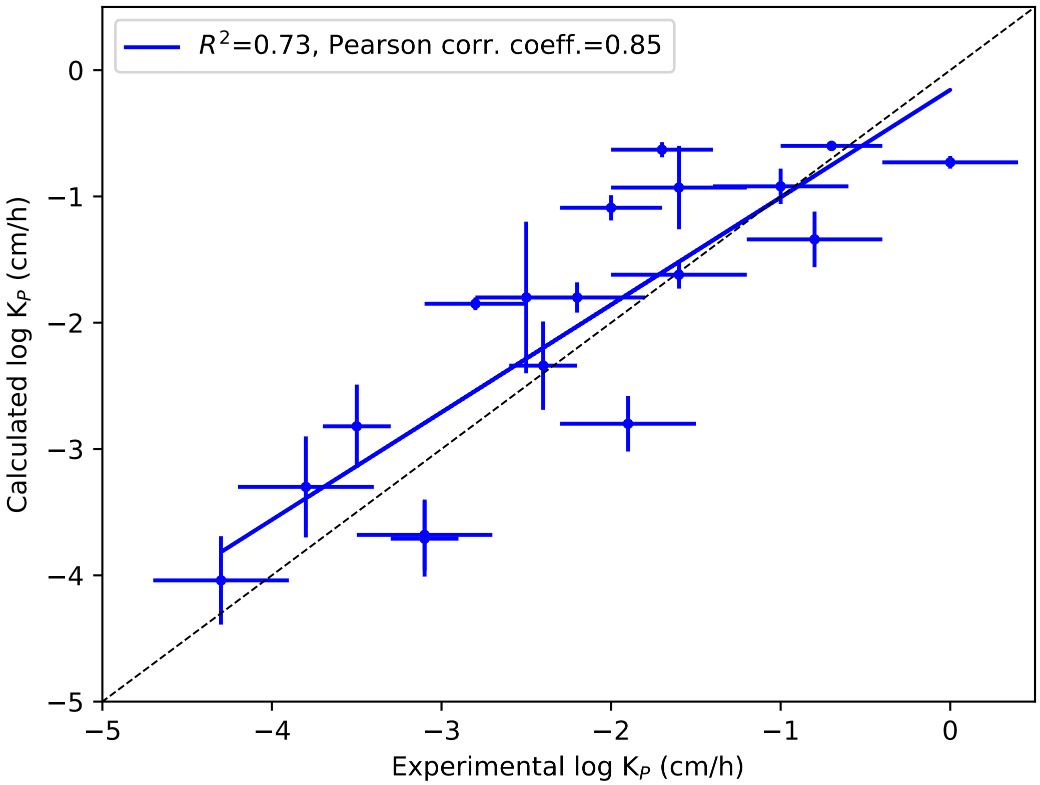


**Fig. S7.** The relationship between experimental log $K_P$ and those calculated from MD simulations, with the two largest outliers (urea and hydromorphone) excluded. The blue line shows the linear regression, whereas the dashed black line is the identity line. The error bars represent 1 SEM (standard error of the mean), which is approximated for experimental values. Details about the approximation is presented in the header of Table S1.

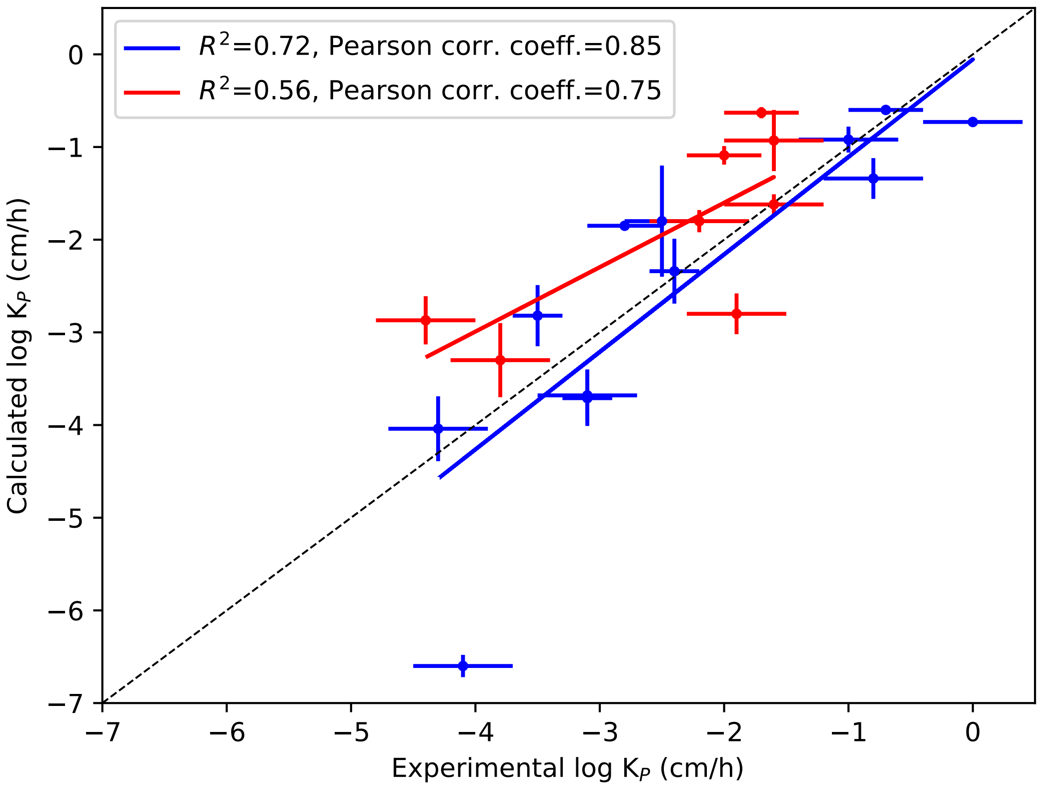


**Fig. S8.** Separating molecules that are not ionizable (shown in blue) under the *ex vivo/in vitro* conditions from those that are ionizable (shown in red). The ionizable compounds are codeine, diclofenac, hydromorphone, ibuprofen, lidocaine, naproxen, nicotine and salicylic acid.

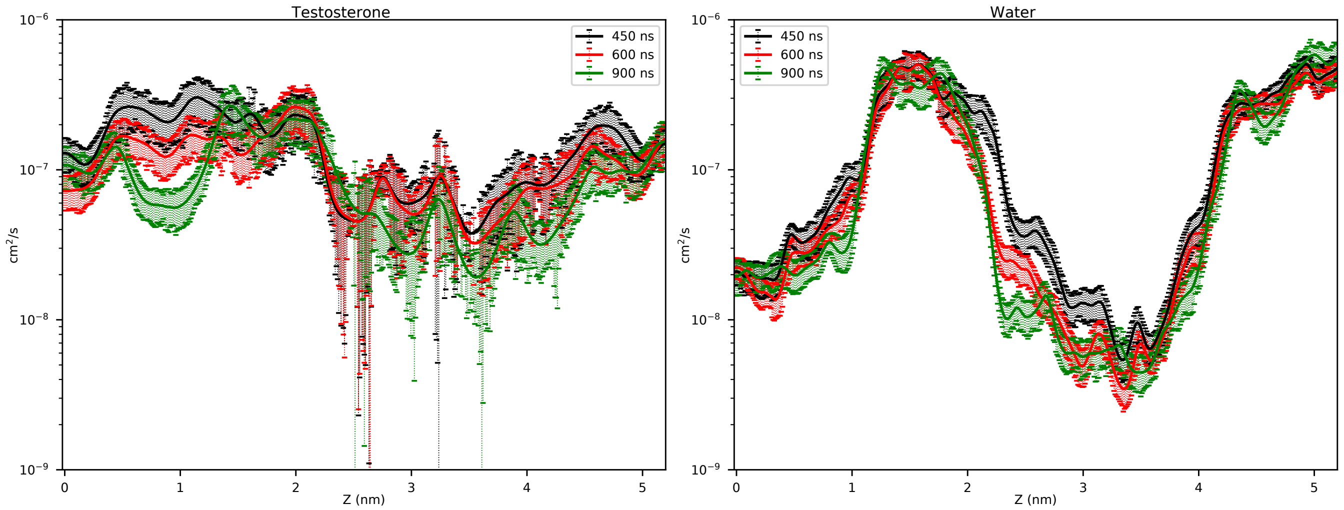


**Fig. S9.** Calculated diffusion coefficients of testosterone and water from extended AWH simulations (24 communicating AWH walkers). The average diffusion coefficients through the system for testosterone were $1.4 \times 10^{-7}\,\mathrm{cm^2\,s^{-1}}$, $1.1 \times 10^{-7}\,\mathrm{cm^2\,s^{-1}}$ and $9.5 \times 10^{-8}\,\mathrm{cm^2\,s^{-1}}$ at $450\,\mathrm{ns}$, $600\,\mathrm{ns}$ and $900\,\mathrm{ns}$, respectively. For water they were $1.6 \times 10^{-7}\,\mathrm{cm^2\,s^{-1}}$, $1.4 \times 10^{-7}\,\mathrm{cm^2\,s^{-1}}$ and $1.5 \times 10^{-7}\,\mathrm{cm^2\,s^{-1}}$ at the same time points.

**Table S2.** Permeability coefficients of testosterone and water from extended AWH simulations. The AWH uncertainties from one single set of simulations (24 communicating AWH walkers) are rough estimates and are expected to be too low.

| | Simulation time (ns) per AWH walker (24 walkers) | | |
|---|---|---|---|
| **Permeant** | **450** | **600** | **900** |
| Testosterone | $-3.4 \pm 0.3$ | $-3.3 \pm 0.3$ | $-3.2 \pm 0.3$ |
| Water | $-4.1 \pm 0.3$ | $-4.2 \pm 0.2$ | $-4.2 \pm 0.2$ |
